# Separase Protease Activity is Required for Cytokinesis in addition to Chromosome Segregation

**DOI:** 10.1101/069906

**Authors:** Xiaofei Bai, Joshua N. Bembenek

## Abstract

Chromosomal segregation and cytokinesis are tightly regulated processes required for successful cell division. The cysteine protease separase cleaves a subunit of the cohesin complex to allow chromosome segregation at anaphase onset. Separase also regulates meiotic cortical granule exocytosis and vesicle trafficking during cytokinesis, both of which involve RAB-11. Separase has non-proteolytic signaling functions in addition to its role in substrate cleavage, and its mechanism in exocytosis is unknown. We sought to determine whether separase regulates RAB-11 vesicle exocytosis through a proteolytic or non-proteolytic mechanism. To address this question, we generated a protease-dead separase, SEP-1^PD^::GFP, and unexpectedly found that it is dominant negative. Consistent with its role in cohesin cleavage, SEP-1^PD^::GFP causes chromosome segregation defects. Depletion of the substrate subunit of cohesin rescues this defect, suggesting that SEP-1^PD^::GFP impairs cohesin cleavage by a substrate trapping mechanism. We investigated whether SEP-1^PD^::GFP also impairs RAB-11 vesicle trafficking. SEP-1^PD^::GFP causes a low rate of cytokinesis failure that is synergistically exacerbated by depletion of the core exocytic t-SNARE protein SYX-4. Interestingly, SEP-1^PD^::GFP causes an accumulation of RAB-11 vesicles at the cleavage furrow site and delayed the exocytosis of cortical granules during anaphase I. Depletion of *syx-4* further enhanced RAB-11::mCherry and SEP-1^PD^::GFP plasma membrane accumulation during cytokinesis. These findings suggest that the protease activity of separase is required for the exocytosis of RAB-11 vesicles during cortical granule exocytosis and mitotic cytokinesis.

**Author Summary:** The defining event of cell division is the equal distribution of the genetic material to daughter cells. Once sister chromatids align on the metaphase plate, the cell releases the brakes to enter anaphase by activating the protease separase. Separase cleaves the cohesin glue holding duplicated sister chromatids together allowing chromosome segregation. Subsequently, the cell must orchestrate a complex series of anaphase events to equally partition the chromatids and the rest of the cellular components into two distinct daughter cells during cytokinesis. Separase has multiple functions during anaphase to help regulate several key events, including promoting vesicle exocytosis required for cytokinesis. Previous studies have shown that separase can exert control over different events either through substrate cleavage, or by triggering signaling pathways. Here we analyze the cellular functions of separase that are impacted by protease inactive separase. Our results show that separase cleaves cohesin to promote chromosome segregation and also cleaves another independent substrate to promote exocytosis. These findings provide a foundation for understanding the molecular control of separase in exocytosis and indicate that separase has multiple independent substrates that it must cleave to execute various functions. This mechanism may enable the cell to coordinate multiple anaphase events with chromosome segregation.

## Introduction

Faithful cell division depends on coordinated regulation of chromosome segregation and cytokinesis. Chromosome segregation requires equal partitioning of sister chromatids that are duplicated and linked together by cohesin during mitotic S-phase [1]. At the onset of anaphase, the kleisin subunit of cohesin, SCC-1, is cleaved by the caspase-like cysteine protease separase, allowing sister chromatid separation [2]. Separase is a large protease with two sub-domains, the pseudo-protease domain (PPD) and active protease domain (APD) as well as an extended ARM repeat region in the N-terminus [3, 4]. The canonical role of separase is to cleave SCC-1, which allows chromosome segregation during mitotic and meiotic anaphase in all eukaryotic organisms studied to date [5]. The proteolytic function of separase is required for several other cell cycle events in anaphase. In budding yeast, separase cleaves the kinetochore and spindle associated protein Slk19, which stabilizes the anaphase spindle [6, 7]. Additionally, separase cleaves the pericentriolar material proteins kendrin and pericentrin B to regulate centriole licensing in mammalian cells [8, 9]. Interestingly, separase cleaves itself at multiple adjacent sites [10]. The auto-cleaved fragments still maintain catalytic activity, and self-cleavage plays important roles in controlling cell cycle progression, separase activity and chromosome segregation [11, 12]. These proteolytic functions stress the importance of identifying the distinct roles of separase and its substrates in both meiosis and mitosis.

In addition to its roles as a protease, several non-proteolytic functions of separase have been identified. At anaphase onset, separase-dependent activation of the Cdc14 early anaphase release (FEAR) pathway initiates mitotic exit in budding yeast [13]. A protease dead separase mutant is still sufficient to initiate mitotic exit but cannot promote cohesin cleavage and spindle elongation [14]. Interestingly, Cdc14 has been shown to promote cytokinesis by regulating ER to bud neck trafficking of chitin synthase and directly dephosphorylating several bud neck targets [15-19]. Separase is also known to bind and inhibit CDK-1 in mammalian cells through an unstructured region between the catalytic and N-terminal domain [20-22]. Consistent with this, several studies have shown that expression of catalytically inactive separase can rescue multiple aspects of separase function [21, 23]. In oocytes, expression of inactive separase can rescue polar body extrusion, a highly asymmetric form of cytokinesis, after knockdown of endogenous separase [23]. These earlier studies would suggest the hypothesis that protease dead separase might be capable of promoting the cytokinetic functions of separase. However, our unexpected observation that protease dead separase is dominant negative in *C. elegans* suggests that it interferes with endogenous separase function. This provides a novel opportunity to investigate the cellular functions that require the protease activity of separase, and to investigate how separase regulates cytokinesis [24].

*Caenorhabditis elegans* is a powerful model system for addressing fundamental cell cycle events. Oocytes mature and undergo fertilization every 25 minutes, then complete meiosis and initiate the mitotic cell divisions within an hour in utero, all of which can be imaged with relative ease [25]. In *C. elegans*, separase performs multiple functions during the oocyte-to-embryo transition in the first meiotic division and the mitotic metaphase-to-anaphase transition. Separase is essential for homologous chromosome disjunction through cleaving meiosis-specific kleisin subunit Rec8 [26]. During anaphase I, separase cleaves the CENP-A related protein, CPAR-1, which may regulate the metaphase-anaphase transition in *C. elegans* [27]. Separase is involved in centriole disengagement during male spermatocyte meiosis [28] and regulates the separation and duplication of sperm-derived centrioles in embryos at the meiosis-mitosis transition [29]. During mitosis, separase cleaves the mitotic cohesin kleisin subunit SCC-1 to promote
chromosome segregation [26, 30]. Whether *C. elegans* separase has the same conserved non-proteolytic functions such as CDK-1 inhibition is unknown, as is whether other protease dead separase mutants are dominant negative in other systems.

Our previous studies have defined an essential function for separase in the regulation of vesicle exocytosis during anaphase. Separase inactivation causes eggshell defects and cytokinesis failures, both of which are due to defects in vesicle trafficking. During anaphase I, separase localizes to cortical granules and is required for their exocytosis, which is necessary for eggshell formation [31]. Simultaneously, it localizes to the base of the polar body and is required for successful cytokinesis during polar body extrusion (PBE). RAB-11, a small GTPase that regulates trafficking at recycling endosome and is essential for cytokinesis in several systems, is also found on cortical granules and the base of the polar body and is required for both events in anaphase I [32]. Further study indicated that separase is also required for cytokinesis during mitosis [33]. Interestingly, depletion of separase in *C. elegans* with RNAi enhanced the accumulation of RAB-11 positive vesicles at the ingressing furrow and midbody, suggesting a role of separase in exocytosis during cytokinesis [33]. Furthermore, the role of separase in exocytosis is independent of its function in chromosome segregation as a unique hypomorphic mutant that maps to the N-terminal domain promotes mostly normal chromosome segregation, while CGE and cytokinesis still remain severely affected [31, 33]. These studies demonstrate that CGE is under the control of the same cellular machinery that regulates membrane trafficking during polar body extrusion and mitotic cytokinesis. Separase has been also found in plant and mammalian systems to regulate membrane trafficking [34, 35], suggesting that separase may have a conserved function in regulating membrane trafficking.

There are many open questions about the exact mechanism of how separase regulates RAB-11 vesicle exocytosis. Previous studies in mouse oocytes suggest that separase has a non-proteolytic role in polar body extrusion, and thus possibly in vesicle trafficking [23]. However, we recently reported the unexpected observation that SEP-1^PD^::GFP is dominant negative in *C. elegans* [24]. Here, we investigated cellular phenotypes to understand what processes are impaired by SEP-1^PD^::GFP in *C. elegans* and whether vesicle trafficking is affected. We used high-resolution confocal microscopy to observe SEP-1^PD^::GFP phenotypes during meiosis I and mitotic cytokinesis. We show that SEP-1^PD^::GFP impairs both chromosome segregation and RAB-11 vesicle trafficking. Depletion of the substrate, cohesin *scc-1*, substantially rescues lagging and bridging anaphase chromosomes in SEP-1^PD^::GFP embryos, suggesting that SEP-1^PD^::GFP prevents substrate cleavage. SEP-1^PD^::GFP also impairs vesicle exocytosis and genetically interacts with vesicle fusion machinery. Therefore, separase may also cleave a substrate to promote exocytosis during CGE and cytokinesis.

## Results

### SEP-1^PD^::GFP inhibits chromosome separation

To investigate the proteolytic functions of separase in *C. elegans*, we used the *pie-1* promoter for germline expression of a protease-dead separase (C1040S) fused to GFP (SEP-1^PD^::GFP) [24, 33]. We have devised two methods to propagate animals carrying the dominant negative protease dead separase and have applied them to characterize the phenotype caused by stable expression of protease-dead separase [24]. Depending on the experimental setup and desired genotype, our conditions lead to SEP-1^PD^::GFP expression from either one or two copies of the transgene in a wild type background with endogenous separase expression. We characterized multiple independently generated SEP-1^PD^::GFP transgenic lines obtained by microparticle bombardment to identify the most reproducibly behaved lines. Two lines (WH520 and WH524) behave as chromosomal-integrated alleles with consistent expression of the protease-dead separase that lead to consistent phenotypes, while other lines were less consistent (S1 Fig). We used WH520 to characterize cellular phenotypes, which has nearly 100% embryo lethality after 5 generations off GFP RNAi (which we will call homozygous SEP-1^PD^::GFP) and about 70% lethality in F2 embryos using the backcross propagation strategy (labeled as SEP-1^PD^::GFP/-) (S1 Fig). In contrast, expression of SEP-1^WT^::GFP causes no lethality and can fully rescue mutant separase embryos [24, 33]. Therefore, expression of SEP-1^PD^::GFP in the wild type background with endogenous separase consistently causes embryo lethality.

Separase is well known to cleave cohesin to allow chromosome segregation. We hypothesized that SEP-1^PD^::GFP is dominant negative in part because it may bind cohesin but would be unable to cleave it, thus preventing endogenous separase from cleaving cohesin and inhibiting chromosome separation. Separase has several conserved substrates that are found in *C. elegans* and mammalian cells, including cohesin. Prior to anaphase onset, SEP-1^WT^::GFP and SEP-1^PD^::GFP show identical localization patterns and both show equivalent localization to chromosomes [33]. However, in anaphase, when separase becomes catalytically active and would bind to substrates, SEP-1^PD^::GFP displays ectopic localization at centrioles and the central spindle where known substrates are cleaved by separase in other system [33], consistent with the hypothesis that it has enhanced association with substrates. In order to investigate the effects of SEP-1^PD^::GFP on chromosome segregation, we compared embryos expressing H2B::mCherry to label the chromosome and homozygous SEP-1^PD^::GFP or SEP-1^WT^::GFP. We defined anaphase onset as the time point when the width of the chromosome signal increases due to spindle forces pulling sister chromatids apart, which always occurs very quickly after chromosome alignment on the metaphase plate in both SEP-1^PD^::GFP and SEP-1^WT^::GFP. Consistent with our hypothesis, chromosome segregation during first anaphase was significantly delayed in homozygous SEP-1^PD^::GFP compared to SEP-1^WT^::GFP embryos (Fig 1 A-L). To ensure cell cycle timing was normal, we quantified the time from nuclear envelop breakdown (NEBD) to furrow ingression in homozygous SEP-1^PD^::GFP embryos and did not observe a significant delay of global cell cycle events as compared with SEP-1^WT^::GFP (S2 Fig).

**Fig 1.**
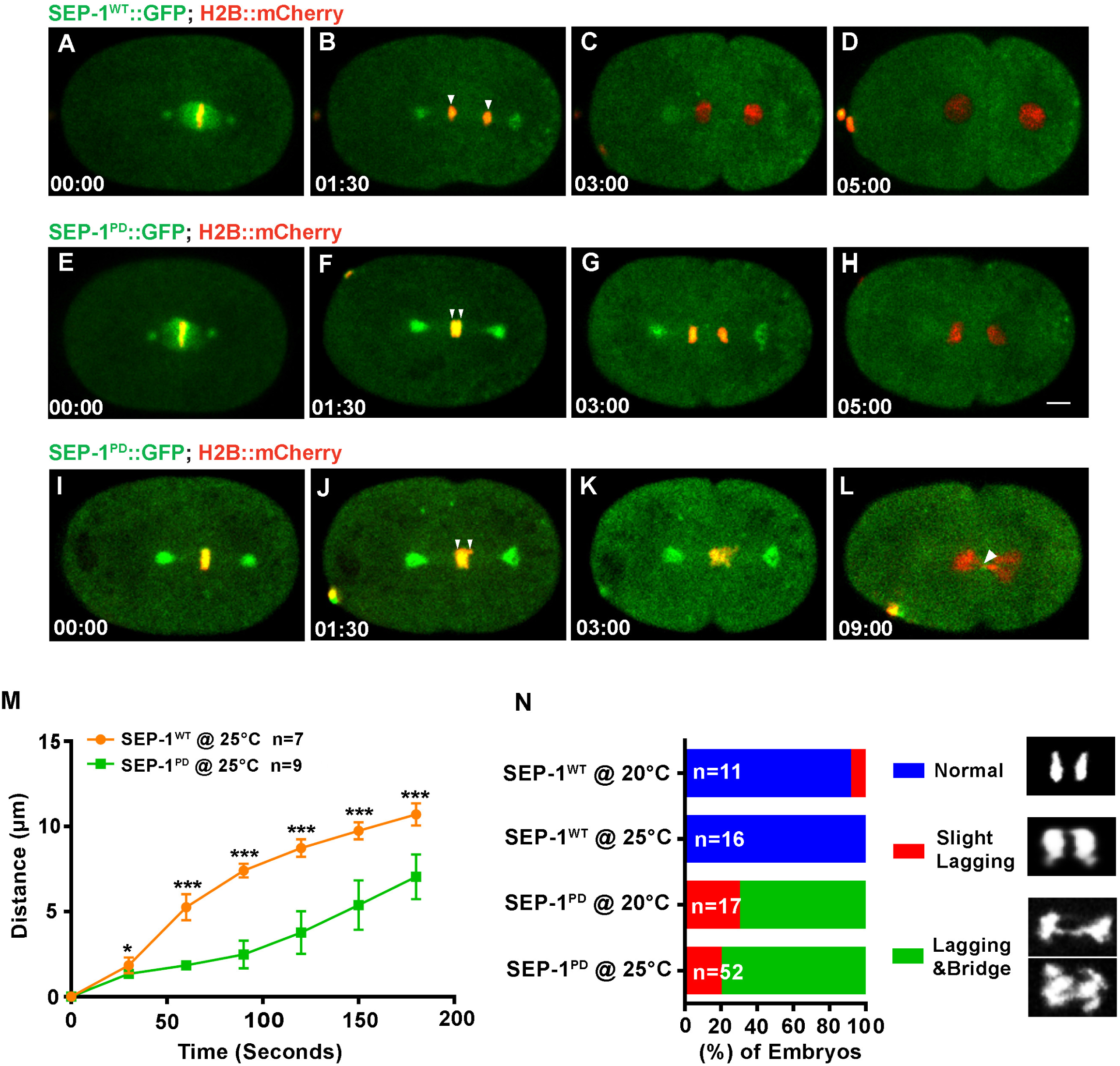
SEP-1^PD^::GFP causes chromosome segregation defects during mitosis. Representative images of mitotic chromosome segregation in SEP-1^WT^::GFP expressing embryos (A-D, green) or homozygous SEP-1^PD^::GFP (green) embryos with slightly lagging (E-H) and bridging (I-L) co-expressing H2B::mCherry (red). M. Average distance between separating sister chromatids during anaphase in SEP-1^WT^::GFP (n=7) or SEP-1^PD^::GFP (n=9) embryos from metaphase to late cytokinesis. N. Percentage of embryos displaying normal chromosome separation (blue), slight lagging chromosomes (red) or chromosome bridges (green) during the first mitosis in embryos expressing either SEP-1^WT^::GFP or SEP-1^PD^::GFP at the temperature indicated (n= number of embryos imaged). Insert shows H2B::mCherry images scored as normal, slight lagging and bridging. Scale Bars, 10 μm. P-values: * =<0.05; ***=<0.0001; ns= not significant.

Quantification of the distance that chromosomes separate after anaphase onset showed on average a 3.7 micron lag in separation over time in embryos expressing homozygous SEP-1^PD^::GFP (Fig 1 M). Homozygous SEP-1^PD^::GFP embryos had some variation in the severity of segregation defects, from slightly lagging ultrafine bridges (in 10/52 SEP-1^PD^::GFP embryos at 25 °C) to more severe lagging and bridging chromosomes (in 42/52 homozygous SEP-1^PD^::GFP embryos at 25 °C) (Fig 1N), which was virtually absent from WT (in 0/16 SEP-1^WT^::GFP embryos at 25 °C). The delayed chromosome separation was more severe at 25 °C than at 20 °C, which is likely due to higher transgene expression (Fig 1N & Movie S1). We also investigated chromosome segregation during anaphase I. Interestingly, homozygous SEP-1^PD^::GFP embryos also displayed chromosome segregation defects during meiotic anaphase (Fig 2 A-H). We measured the delay in separation over time and observed a less severe but significant delay in chromosome segregation (Fig 2 I). In addition, the bridging defects were not as severe as observed in mitosis (slightly lagging bridge observed in 0/8 SEP-1^WT^::GFP embryos at 25 °C; in 7/15 homozygous SEP-1^PD^::GFP at 25 °C, Fig 2J & Movie S2). These data indicate that homozygous SEP-1^PD^::GFP impairs chromosome segregation during both meiosis and mitosis, likely due to impaired cohesin cleavage.

**Fig 2.**
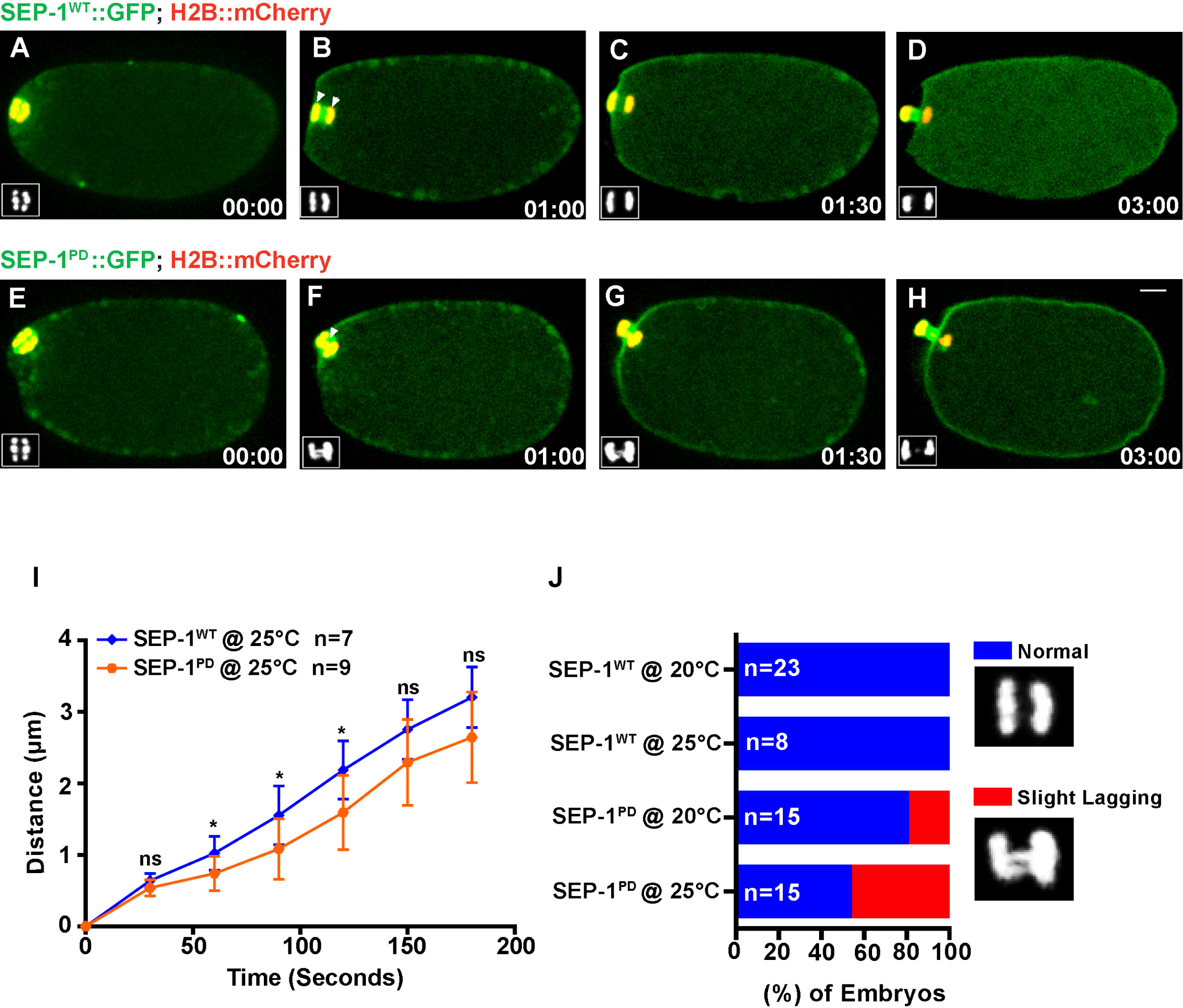
SEP-1^PD^::GFP causes chromosome segregation defects during meiosis I. Representative images of meiotic chromosome segregation in SEP-1^WT^::GFP (A-D, green) or homozygous SEP-1^PD^::GFP expressing embryos (E-H, green) co-expressing H2B::mCherry (red). Lower left insets show H2B::mCherry. I. Average distance between chromosomes during anaphase in SEP-1^WT^::GFP or homozygous SEP-1^PD^::GFP. J. Percentage of embryos displaying normal chromosome separation (blue), lagging chromosomes (red) during the anaphase I in embryos expressing either SEP-1^WT^::GFP or homozygous SEP-1^PD^::GFP (n= number of embryos imaged). Insets show examples scored as normal or lagging chromosomes during anaphase I. Scale Bars, 10 μm. P-values: * =<0.05; ns= not significant.

If our hypothesis that cohesin cleavage is impaired by SEP-1^PD^::GFP is correct, we would expect that partial depletion of *scc-1* by RNAi would alleviate the chromosome segregation defects. We carefully titrated the degree of RNAi depletion (feeding RNAi 24 hours at 20°C and 25 °C) to achieve a mild level of *scc-1* depletion to avoid causing severe chromosome segregation defects due to loss of cohesin [30]. At both 20°C and 25 °C, *scc-1* RNAi causes only mild lethality in WT (Fig 3A, B) but significantly rescues the homozygous SEP-1^PD^::GFP embryonic lethality from 100% down to 22% ± 10.34 at 25 °C (Fig 3B). Chromosome segregation defects were significantly alleviated after depletion of *scc-1* (RNAi) in homozygous SEP-1^PD^::GFP embryos (Fig 3 C-F). Homozygous SEP-1^PD^::GFP depleted of *scc-1* also had normal kinetics of chromosome segregation in anaphase (Fig 3G) and much less severe bridging defects (28/42 normal, 9/42 slight lagging, 5/42 severe bridges, Fig 3H). Therefore, reducing the amount of cohesin largely rescues the chromosome segregation defects caused by expressing SEP-1^PD^::GFP together with endogenous separase in *C. elegans*. Presumably this is because there is less substrate that must be cleaved, reducing the amount of cohesin that endogenous separase must cleave in the presence of SEP-1^PD^::GFP to allow chromosome segregation. These observations are consistent with our hypothesis that SEP-1^PD^::GFP inhibits substrate cleavage, causing a dominant phenotype.

**Fig 3.**
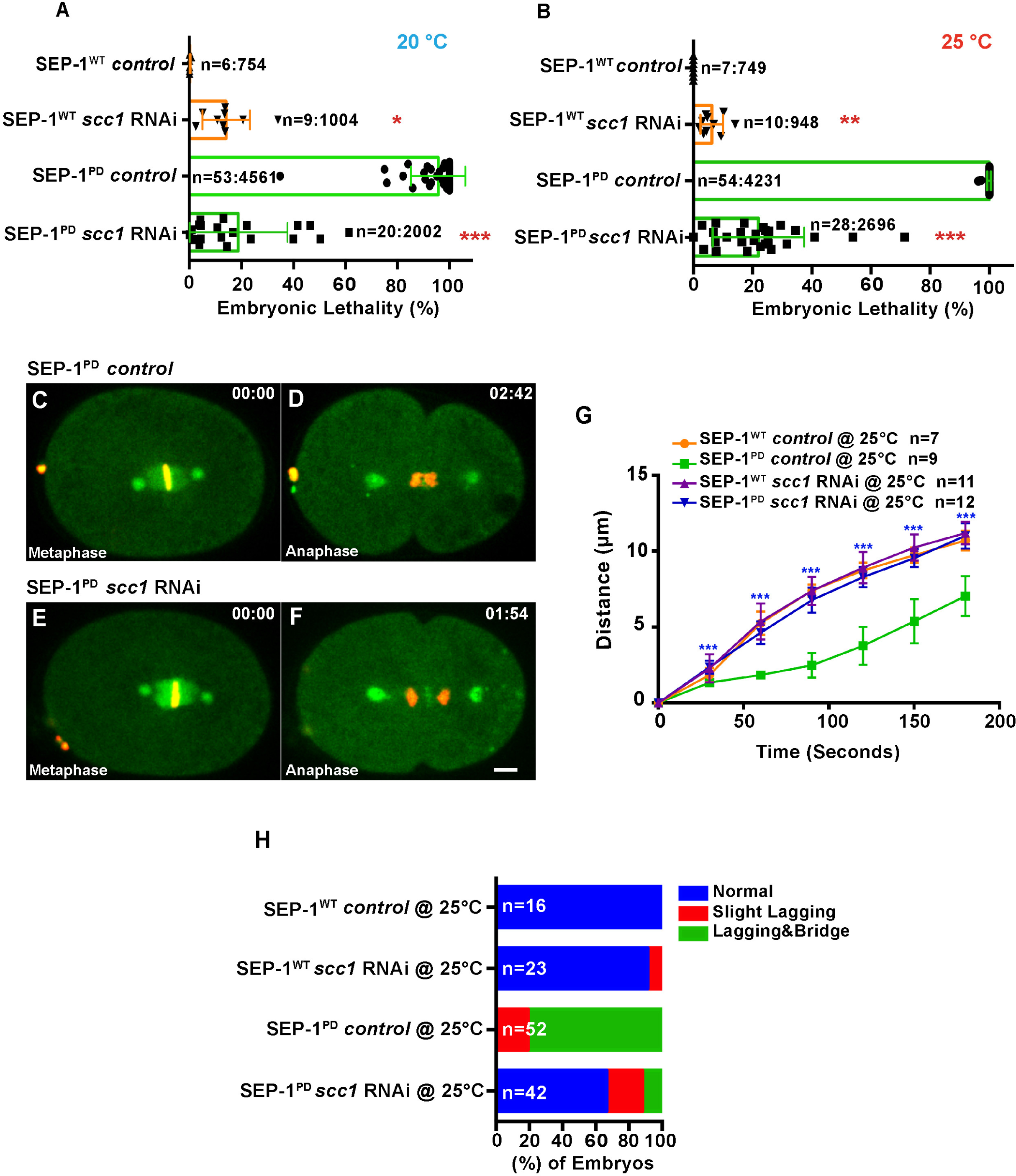
Cohesin depletion rescues chromosome segregation defects caused by SEP-1^PD^::GFP. (A, B) Partial cohesin depletion significantly rescues the SEP-1^PD^::GFP embryonic lethality at both 20°C and 25 °C. (C-F) Chromosome segregation defects were significantly alleviated after partial depletion of *scc-1* in homozygous SEP-1^PD^::GFP (green) embryos (DNA in red). G. Distance between separating sister chromatids during anaphase in SEP-1^WT^::GFP or SEP-1^PD^::GFP control or with *scc-1 (RNAi)*. (H) Percentage of embryos displaying normal chromosome separation (blue), lagging chromosomes (red) or chromosome bridges (green) during the first mitosis in embryos expressing SEP-1^WT^::GFP or SEP-1^PD^::GFP with and without *scc-1 (RNAi)* treatment (n= number of embryos imaged). Scale Bars, 10 μm. P-values: * =<0.05; **=<0.01; ***=<0.0001.

### SEP-1^PD^::GFP expression causes cytokinesis failures independent of cohesin

In addition to the canonical function of separase in chromosome segregation, separase is required for cytokinesis by regulating vesicle exocytosis [33]. If separase has a substrate that it must cleave in order to promote vesicle exocytosis during cytokinesis, we postulated that SEP-1^PD^::GFP would inhibit this process similar to the way it impairs chromosome segregation. We tested whether homozygous SEP-1^PD^::GFP embryos fail cytokinesis using live imaging. Interestingly, we found some homozygous SEP-1^PD^::GFP embryos with multipolar spindles, indicative of cytokinesis failure, in one cell through two cell stages (in 2/52 homozygous SEP-1^PD^::GFP embryos; in 0/17 SEP-1^WT^::GFP embryo; in 0/13 N2 at 25 °C. Fig 4A). Additionally, cytokinesis failures are sporadic and are often seen in older embryos, but are difficult to quantify because cells that fail cytokinesis often undergo multipolar division and cellularize (data not shown). These data indicate that SEP-1^PD^::GFP expression does cause cytokinesis failure, suggesting that there may be a substrate cleavage event required for cytokinesis.

**Fig 4.**
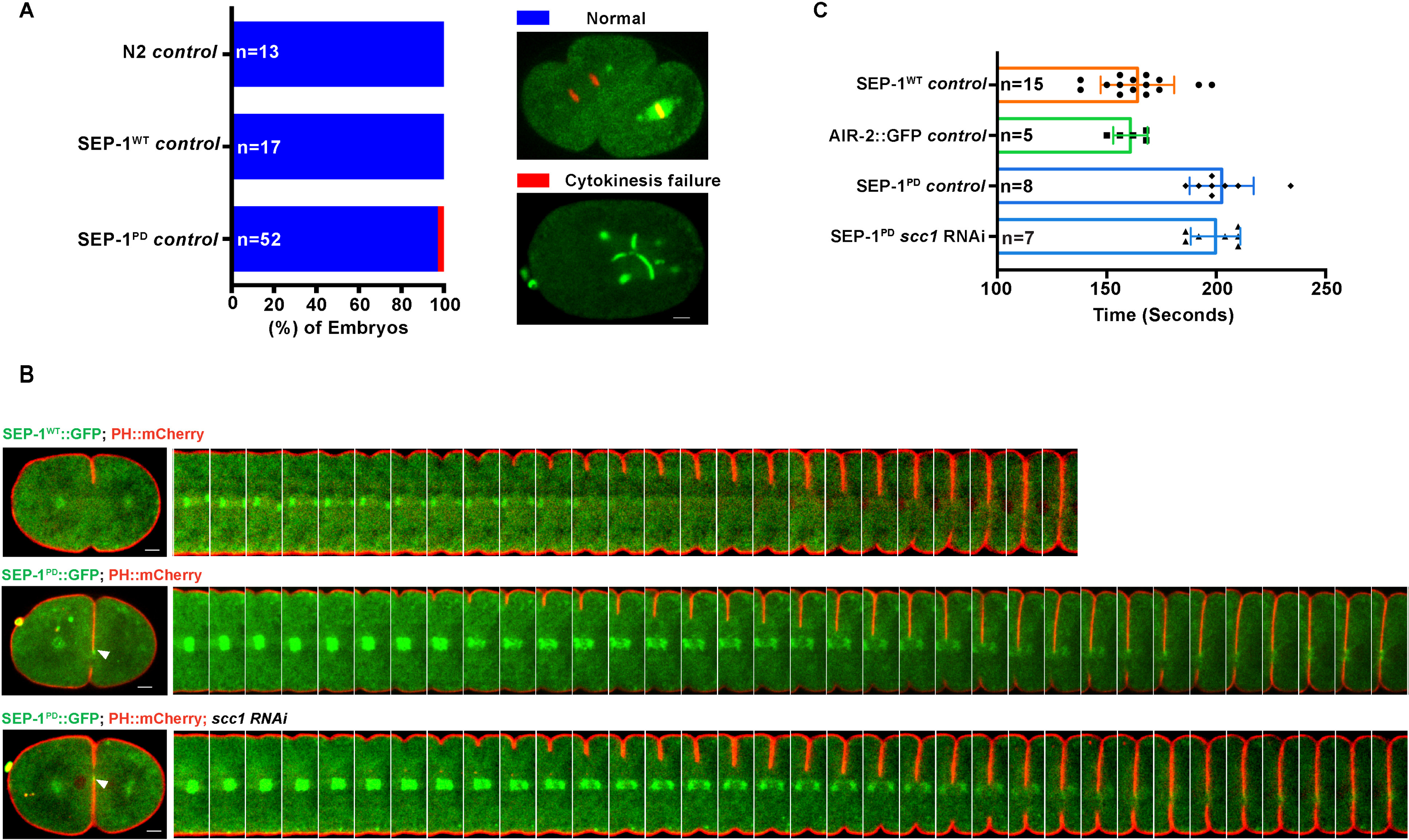
SEP-1^PD^::GFP causes cytokinesis defects. (A) Percentage of embryos displaying normal cell division (blue) and cytokinesis failure (red) during first mitotic division in N2 wild type, SEP-1^WT^::GFP or homozygous SEP-1^PD^::GFP. Right panels show examples scored as normal or cytokinesis failure. (B) Kymograph of the furrow region shows PH::mCherry (red) in SEP-1^WT^::GFP, homozygous SEP-1^PD^::GFP and SEP-1^PD^::GFP; *scc-1 (RNAi)* during cytokinesis (separase in green). (C) Quantification of the furrow ingression time in SEP-1^WT^::GFP, AIR-2::GFP or homozygous SEP-1^PD^::GFP with and without *scc-1(RNAi)* (n= number of embryos imaged). Scale Bars, 10 μm.

Next, we analyzed the rate of furrow ingression to determine if there are additional defects during cytokinesis despite the low rate of cytokinesis failure. We generated homozygous SEP-1^PD^::GFP and SEP-1^WT^::GFP lines expressing PH::mCherry to observe the plasma membrane during cytokinesis. We imaged furrow ingression in a single focal plane of the central spindle and midbody. SEP-1^PD^::GFP, but not SEP-1^WT^::GFP, is often colocalized with the plasma membrane during furrowing and remains at the midbody for an extended period of time, which could reflect enhanced association with a membrane substrate (Fig 4B). We found that furrow ingression rate in homozygous SEP-1^PD^::GFP embryos was consistently slower compared with the SEP-1^WT^::GFP and AIR-2::GFP control (0.138 μm/second ± 0.003 in homozygous SEP-1^PD^::GFP; 0.169 μm/second ± 0.006 in SEP-1^WT^::GFP; S3 Fig B). We also measured the time from the initiation of furrow ingression until it completed, generating a smooth cell boundary. In SEP-1^WT^::GFP cells, this process took (164 seconds ± 4.3, n=15, Fig 4C). Since SEP-1^WT^::GFP does not label the midbody, we also imaged the midbody maker AIR-2::GFP together with PH::mCherry and found that our measurement of furrow completion timing was accurate (160.8 seconds ± 3.5, n=5, Fig 4C, S3 Fig A). In homozygous SEP-1^PD^::GFP embryos, cytokinesis completion was significantly delayed relative to wild type embryos (202.5 seconds ± 5.2; n=8, Fig 4 C). Therefore, expression of dominant negative SEP-1^PD^::GFP specifically impairs furrow ingression and completion of cytokinesis.

Next, we investigated whether the role of separase during cytokinesis is related to cohesin cleavage or whether it depends on an independent pathway. Cohesin is the critical target of separase in chromosome segregation and is also found on the centrosome where it is cleaved during centriole licensing [36]. A function for cohesin during cytokinesis has not been previously reported. If the dominant negative phenotype of SEP-1^PD^::GFP during cytokinesis depends on defects in cohesin cleavage, we expected that depletion of cohesin would rescue furrowing defects in addition to rescuing chromosome segregation. In order to examine this, we examined furrowing in SEP-1^PD^::GFP embryos depleted of cohesin. Interestingly, partial depletion of *scc-1* did not rescue the delay of furrow closure (199.7 seconds ± 4.3, n=7, Fig 4C) or the furrow ingression rate in homozygous SEP-1^PD^::GFP embryos (0.142 μm/second ± 0.003 in homozygous SEP-1^PD^::GFP *scc-1* RNAi; n=7, S3 Fig B), even though chromosome segregation is largely rescued (Fig 3). This indicates that the cytokinesis defects caused by SEP-1^PD^::GFP are independent of the chromosome segregation defects. In addition, this result suggests that separase has another substrate besides cohesin that it cleaves in order to promote cytokinesis.

### SEP-1^PD^::GFP genetically interacts with essential exocytosis machinery

Given that separase regulates cytokinesis by promoting RAB-11 vesicle exocytosis, we investigated whether SEP-1^PD^::GFP interferes with exocytosis. We first tested whether there was a genetic interaction between SEP-1^PD^::GFP and the t-SNARE *syx-4*. SYX-4 is a core part of the exocytosis fusion machinery and is localized to the plasma membrane where it is required for cytokinesis in *C. elegans* [37]. Therefore, we expected that combining SEP-1^PD^::GFP expression and depletion of *syx-4* would greatly exacerbate the cytokinesis failure rate if they both inhibit exocytosis. *syx-4* RNAi is inefficient and causes highly variable phenotypes compared with other genes [37]. We carefully calibrated RNAi treatment and determined that 30-36 hours feeding *syx-4* RNAi was an optimal intermediate condition, which caused minimal eggshell permeability and cytokinesis defects in wild type embryos. Consistent with our hypothesis, 30-36 hours feeding *syx-4* RNAi synergistically enhanced embryonic cytokinesis defects in embryos expressing homozygous SEP-1^PD^::GFP (in 15/21 cytokinesis failure, Fig 5B, C) as compared with wild type (in 0/23 SEP-1^WT^::GFP embryos; in 2/30 N2 embryos, Fig 5A, C, Movie S3). Therefore, SEP-1^PD^::GFP has a strong negative genetic interaction with *syx-4*, consistent with the hypothesis that they both inhibit exocytosis during cytokinesis.

**Fig 5.**
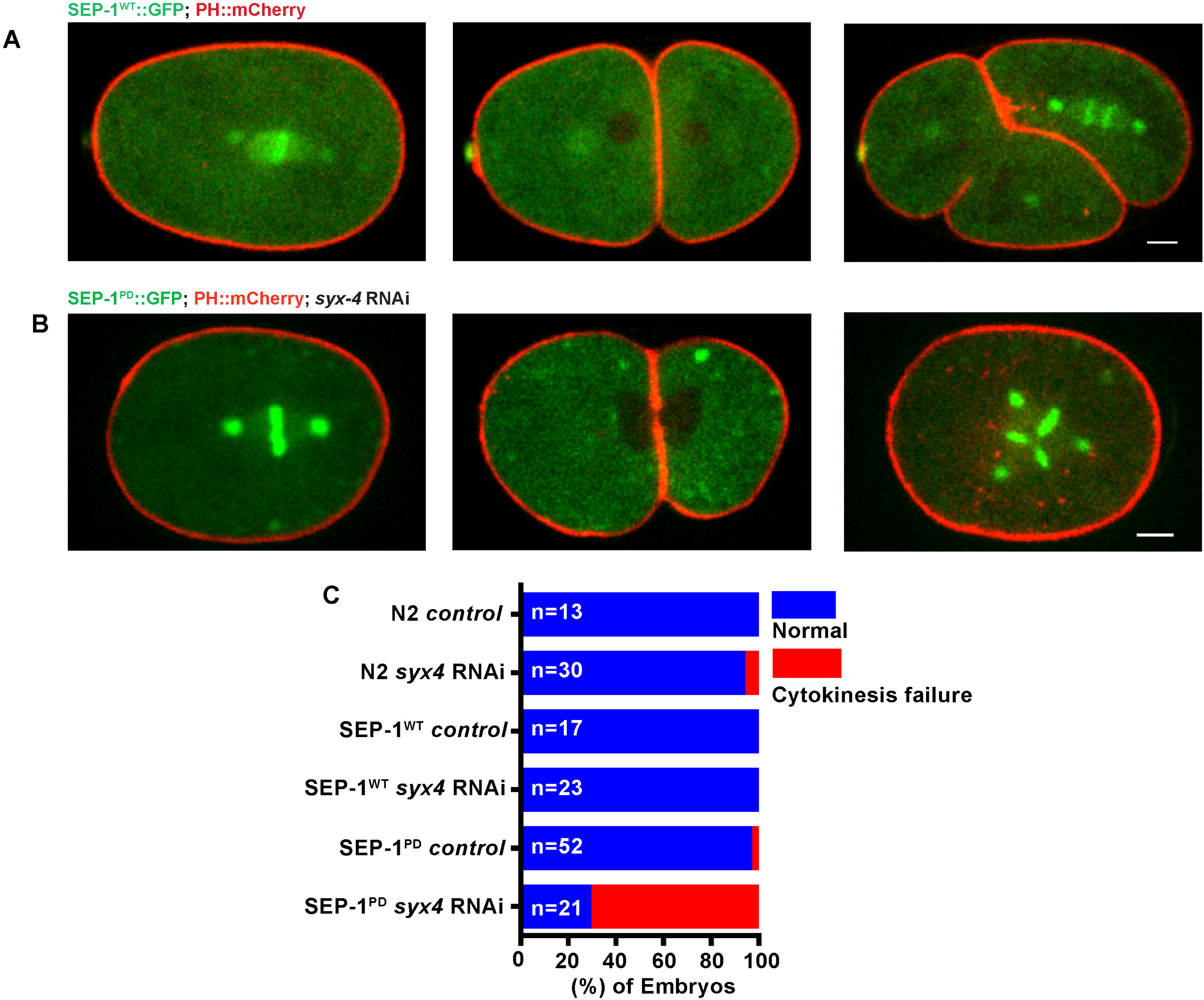
SEP-1^PD^::GFP has a negative genetic interaction with the t-SNARE *syx-4*. (A) Representative images of mitotic cytokinesis in SEP-1^WT^::GFP (A, green) or homozygous SEP-1^PD^::GFP (B, green) embryos co-expressing PH::mCherry (red). (B) Representative images of mitotic cytokinesis failure in homozygous SEP-1^PD^::GFP; PH::mCherry expressing embryos with *syx-4 (RNAi)*, resulting in a one cell embryo with a multi-polar spindle. (C) Percentage of embryos displaying normal cytokinesis (blue) or cytokinesis failure (red) in different conditions (n= number of embryos imaged). Scale Bars, 10 μm.

### SEP-1^PD^::GFP inhibits RAB-11 positive vesicle trafficking during cytokinesis

We next wanted to investigate whether the cytokinesis defects caused by SEP-1^PD^::GFP expression were due to the inhibition of RAB-11 positive vesicle trafficking. Despite several attempts we were unable to generate homozygous SEP-1^PD^::GFP expressing RAB-11::mCherry, indicative of a negative genetic interaction. However, we could generate viable hemizygous SEP-1^PD^::GFP/- and RAB-11::mCherry/-F1 animals that reproducibly expressed both transgenes in order to film F2 embryos. Since the protein in newly fertilized F2 embryos is synthesized by the F1 maternal syncytial germline, each embryo will have the same cytoplasmic expression of SEP-1^PD^::GFP/- and RAB-11::mCherry despite having different genotypes. Although the cytokinesis phenotypes in SEP-1^PD^::GFP/- expressing RAB-11::mCherry are less severe than homozygous SEP-1^PD^::GFP, 30-36 hours feeding of *syx-4* RNAi substantially increased the rate of cytokinesis failures (0/30 *syx-4(RNAi)*; SEP-1^WT^::GFP, 0/15 SEP-1^PD^::GFP/-, 5/23 *syx-4(RNAi)*; SEP-1^PD^::GFP/-, S4 Fig and Movie S4). Mounting embryos on an agar pad or in hanging drop gave the same results after treating *syx-4* RNAi in SEP-1^PD^::GFP compared with SEP-1^WT^::GFP embryos, indicating that indirect effects from mounting were not an issue. Therefore, *syx-4* RNAi strongly exacerbates the cytokinesis defects in both hemizygous and homozygous SEP-1^PD^::GFP embryos, although the cytokinesis phenotypes are weaker in the hemizygous embryos.

Next, we imaged RAB-11 vesicle trafficking during cytokinesis in SEP-1^WT^::GFP and SEP-1^PD^::GFP/- embryos. RAB-11 generates exocytic vesicles from a centrosomal compartment of recycling endosomes, and remains associated with those vesicles as they are transported to and exocytosed at the plasma membrane [38-40]. In SEP-1^WT^::GFP and SEP-1^PD^::GFP/- embryos, RAB-11 is normally distributed at centrosomes and throughout the cytoplasm, indicating that early stages of vesicle trafficking are normal (Fig 6 A, B). Interestingly, we found that the expression of SEP-1^PD^::GFP/- resulted in increased and persistent accumulation of RAB-11 vesicles at the cleavage furrow and midbody compared to SEP-1^WT^::GFP expressing embryos, consistent with a defect in exocytosis at the plasma membrane (Fig 6A, B; Movie S5). The Golgi-associated GTPase, RAB-6, was shown to recruit separase to the cortical granule in *C. elegans* embryos [41]. However, we did not observe the accumulation of RAB-6 at the ingressing furrow or midbody during cytokinesis in SEP-1^PD^::GFP/- expressing embryos (S5 Fig). Therefore, SEP-1^PD^::GFP/- specifically inhibits RAB-11 trafficking during cytokinesis.

**Fig 6.**
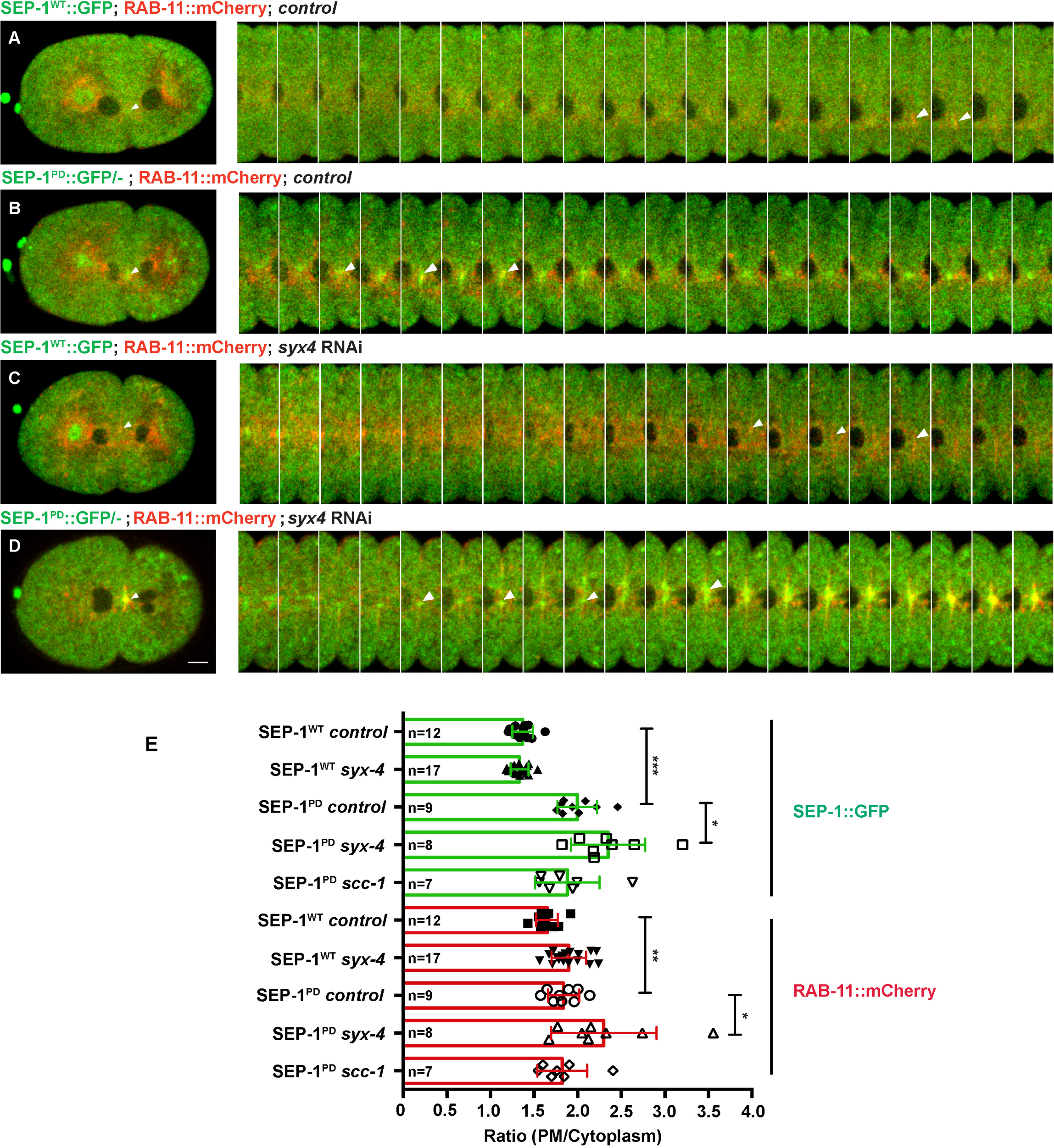
SEP-1^PD^::GFP inhibits RAB-11 positive vesicle trafficking during cytokinesis. (A, B) Representative images and kymograph of RAB-11::mCherry (red) trafficking to the furrow in SEP-1^WT^::GFP (green) or hemizygous SEP-1^PD^::GFP/- (green). Arrowheads denote enhanced RAB-11::mCherry accumulation. (C, D) *syx-4 (RNAi)* enhances RAB-11::mCherry in both SEP-1^WT^::GFP and SEP-1^PD^::GFP/- at the furrow and midbody. (E) Quantification of separase and RAB-11 signals in the midbody during cytokinesis in different conditions (n= number of embryos imaged). Error bars indicated standard error of the mean. P-values: * =<0.05; **=<0.01; ***=<0.0001. Scale Bars, 10 μm

Given that SEP-1^PD^::GFP expression combined with *syx-4* (RNAi) enhances cytokinesis failure (Fig 5C, S4 Fig), we hypothesized that *syx-4* and the proteolytic activity of separase are involved in RAB-11 vesicle exocytosis. To examine this further, we examined whether RAB-11 trafficking was more defective in SEP-1^PD^::GFP/ -; *syx-4* (RNAi) embryos, which might explain the increased cytokinesis failure. We imaged RAB-11 vesicles in embryos expressing both RAB-11::mCherry and SEP-1^PD^::GFP/- with and without 30-36 hours feeding *syx-4* RNAi treatment. Depletion of *syx-4* caused a significantly higher accumulation of both RAB-11::mCherry and SEP-1^PD^::GFP/- at the ingressing furrow and midbody compared with untreated SEP-1^PD^::GFP/- embryos (Fig 6 C, D, E). Unfortunately we could not assay RAB-11 vesicle trafficking under the more severe condition of homozygous SEP-1^PD^::GFP; *syx-4* (RNAi) which has a much higher cytokinesis failure rate, but we expect that RAB-11 accumulation would be even greater. These data are consistent with the hypothesis that separase and RAB-11 are trafficked together on vesicles to the plasma membrane during cytokinesis, and that *syx-4* (RNAi) delays fusion of these vesicles. Finally, we examined whether cohesin would cause any change in RAB-11 vesicle trafficking. Given that partial depletion of *scc-1* does not significantly change the rate of furrow ingression in SEP-1^PD^::GFP embryos (Fig 4B), we expected RAB-11 trafficking would also not be affected. Indeed, depletion of *SCC-1* did not alter the accumulation of RAB-11 vesicles at the furrow in SEP-1^PD^::GFP/- embryos (Fig 6E). These results suggest that separase regulates cytokinesis by hydrolyzing an unknown substrate to regulate exocytosis of RAB-11 positive vesicles.

### SEP-1^PD^::GFP expression delays cortical granule exocytosis

Separase and RAB-11 both localize to cortical granules while SYX-4 localizes to the plasma membrane to promote their exocytosis during meiosis anaphase I [31, 32]. This is an excellent cellular context to investigate exocytosis because separase can be observed directly on these large 1μM vesicles, which release contents required for eggshell formation during anaphase I. We investigated whether SEP-1^PD^::GFP also impairs cortical granule exocytosis (CGE) similar to its effects during cytokinesis. We analyzed whether embryos were permeable to dyes due to disrupted eggshell formation from lack of CGE, but did not observe significant permeability defects. This indicates that SEP-1^PD^::GFP expression does not completely inhibit CGE. To confirm localization, we filmed SEP-1^PD^::GFP/- embryos expressing the cortical granule cargo, CPG-2::mCherry, during anaphase I using hanging drop mounts [31]. We observed that CGP-2::mCherry localizes to cortical granules with both SEP-1^WT^::GFP and SEP-1^PD^::GFP/- as expected (S6 Fig and Movie S6). Interestingly, separase localizes to more vesicles than those labeled by CPG-2::mCherry, indicating that this cargo is only packaged into a subset of cortical granules (S6 Fig, Movie S6). This result is consistent with the heterogeneity of the cortical granule vesicle population observed by transmission electron microscopy [31].

Next, we investigated whether CGE was delayed in homozygous SEP-1^PD^::GFP embryos relative to SEP-1^WT^::GFP. We imaged anaphase I with H2B::mCherry and SEP-1::GFP to observe both chromosomes and cortical granules and quantified the time from anaphase onset until CGE completion during anaphase I. CGE was significantly delayed in homozygous SEP-1^PD^::GFP expressing embryos (198.25 seconds ± 8.0, n=8) compared with SEP-1^WT^::GFP expressing embryos (136.3 seconds ± 5.2, n=7) at 25 °C (Fig 7A-G and Movie S7). In addition, we observed that SEP-1^PD^::GFP remained associated with the plasma membrane for a longer time after CGE (736.4 seconds ± 22.14, n=11) compared with SEP-1^WT^::GFP (220.6 seconds ± 23.6, n=13) (Fig 7H-K & Movie S8). Quantification of the plasma membrane localized signal shows that both SEP-1^WT^::GFP and SEP-1^PD^::GFP initially accumulate on the membrane to similar amounts, but SEP-1^PD^::GFP accumulates to a higher level and remains associated with the membrane for substantially longer (Fig 7L). These data are consistent with the model that SEP-1^PD^::GFP may block cleavage of putative substrate involved in exocytosis and that it may remain bound to a substrate after exocytosis in the plasma membrane.

**Fig 7.**
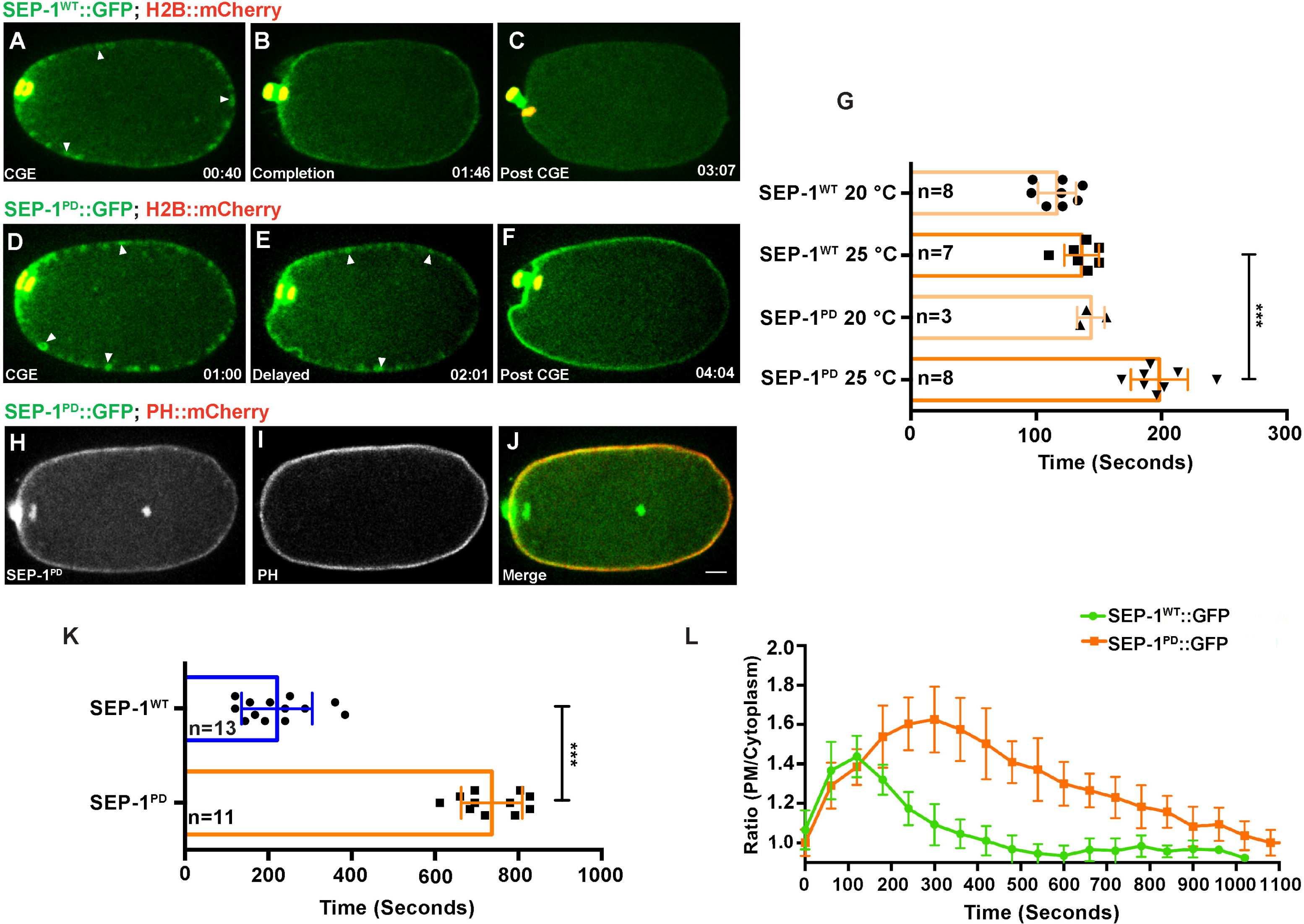
SEP-1^PD^::GFP expression delays cortical granule exocytosis. (A-F) Representative images of separase localization during anaphase I. Localization of SEP-1^WT^::GFP (A, green) and SEP-1^PD^::GFP (D, green) to cortical granules indicated by white arrowheads (H2B::mCherry in red). CGE was delayed in homozygous SEP-1^PD^::GFP (E) compared with SEP-1^WT^::GFP (B) during late anaphase I. (F) SEP-1^PD^::GFP associated with the cortex for a longer time after CGE compared with SEP-1^WT^::GFP (C). (H-J) Colocalization of SEP-1^PD^::GFP (green) with PH::Cherry (red) at the plasma membrane after CGE. (G) Quantification of anaphase onset to completion of CGE. SEP-1^PD^::GFP embryos take longer to finish CGE than SEP-1^WT^::GFP. (K) Average time that SEP-1^WT^::GFP or SEP-1^PD^::GFP remains associated with the plasma membrane after CGE. (L) Ratio of plasma membrane to cytoplasmic SEP-1^PD^::GFP and SEP-1^WT^::GFP after onset of anaphase I. Scale Bars, 10 μm. P-values: ***=<0.0001

### SEP-1^PD^::GFP does not affect RAB-11 after cortical granule exocytosis

RAB-11 localizes to cortical granules and is required for CGE [32]. Therefore, we investigated whether SEP-1^PD^::GFP affects the dynamics of RAB-11 during and after CGE. We filmed meiotic stage embryos expressing SEP-1::GFP/- and RAB-11::mCherry and observed that RAB-11::mCherry localizes to cortical granules several minutes prior to anaphase [32], before either SEP-1^WT^::GFP or SEP-1^PD^::GFP/- localize to cortical granules (Fig 8A, D). Just after anaphase onset, prior to exocytosis, both forms of separase fully co-localize with all RAB-11::mCherry labeled cortical granules prior to exocytosis (Fig 8B, E). Therefore, RAB-11 and separase are localized to the same population of CGs and are sequentially recruited to cortical granules through an orderly process leading to exocytosis in anaphase I (Movie S9). After exocytosis, SEP-1^PD^::GFP/- associated with the plasma membrane for an extended time while RAB-11::mCherry rapidly disappeared (Fig 8F and Movie S9). This result suggests that SEP-1^PD^::GFP/- does not require RAB-11 to remain associated with the plasma membrane, but might bind another unknown substrate. Therefore, RAB-11 and separase may function in parallel but independent pathways to promote exocytosis during anaphase.

**Fig 8.**
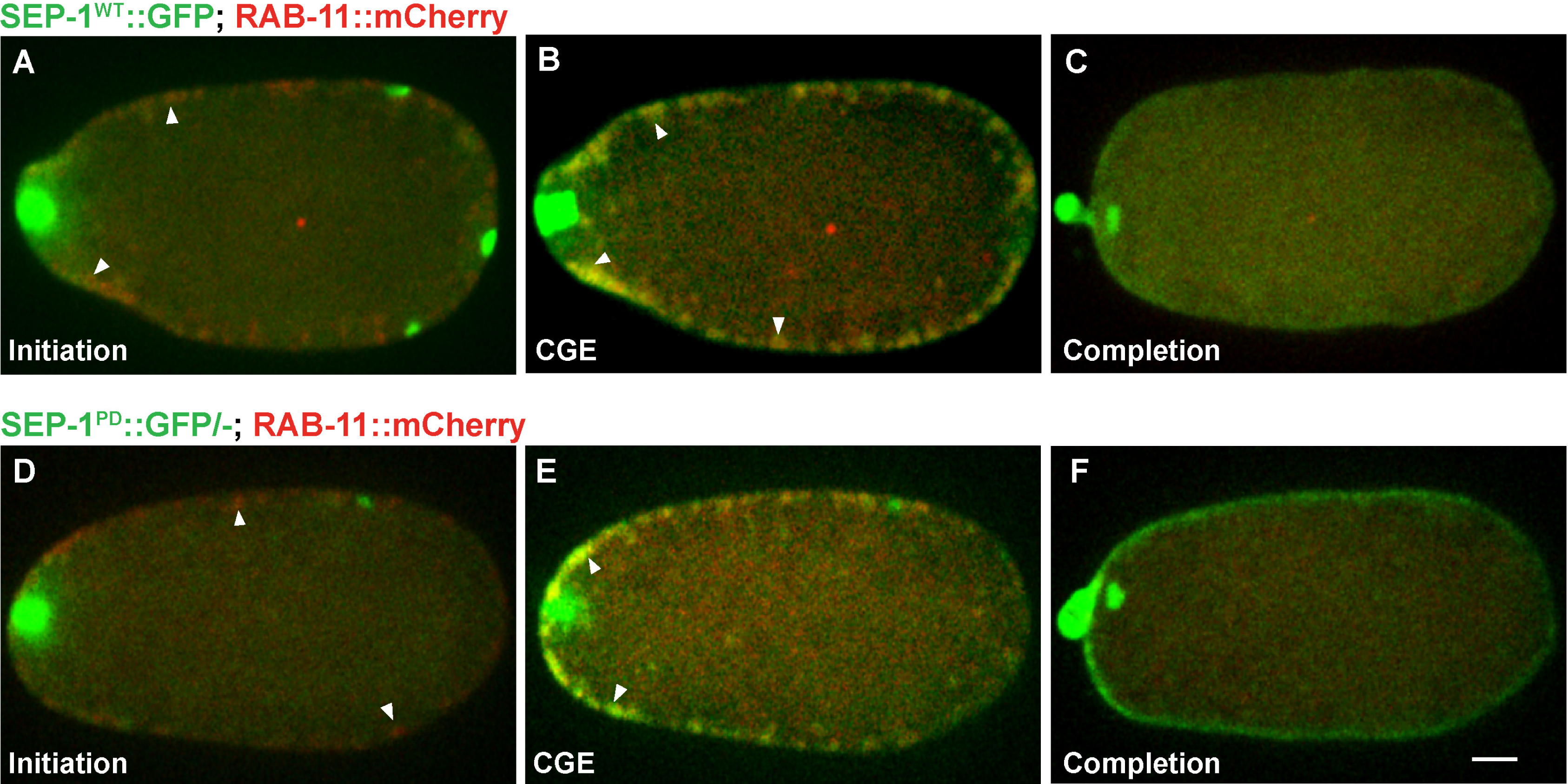
SEP-1^PD^::GFP does not affect RAB-11 after cortical granule exocytosis. Representative images of meiosis I in embryos expressing separase (green) and RAB-11 (red). (A, D) RAB-11 localizes to cortical granules several minutes prior to anaphase, before either SEP-1^WT^::GFP or SEP-1^PD^::GFP/- localize to cortical granules. SEP-1^WT^::GFP (B, green) and SEP-1^PD^::GFP (E, green) colocalize with RAB-11::mCherry (red) on the cortical granules in anaphase I. White arrowheads denote colocalization of separase and RAB-11 on cortical granules. (C, F) After exocytosis, SEP-1^PD^::GFP/- associated with the plasma membrane while SEP-1^WT^::GFP and RAB-11::mCherry rapidly disappeared. Scale Bars, 10 μm.

## Discussion

The mechanism by which separase regulates chromosome segregation is well known, while its function in exocytosis during CGE and cytokinesis needs to be elucidated. Here, we explore whether the proteolytic activity of separase is involved in its membrane trafficking roles. Utilizing our novel observation that protease dead separase is dominant negative, we provide data showing that it interferes with endogenous separase function during chromosome segregation and cytokinesis. Therefore, we hypothesize that separase uses its protease activity to cleave cohesin to allow chromosome segregation and to independently cleave multiple other substrates to promote several events during anaphase, including membrane trafficking during cytokinesis.

During chromosome segregation, the well-established function of separase is to cleave the cohesin subunit SCC-1 during mitosis. Consistent with the hypothesis that SEP-1^PD^::GFP impairs substrate cleavage by the endogenous separase, we observe chromosome segregation defects in SEP-1^PD^::GFP expressing embryos. Furthermore, depletion of SCC-1 substantially recues mitotic chromosome segregation and embryo lethality caused by SEP-1^PD^::GFP. Previously, SCC-1 was not detected on chromosomes after prophase, suggesting that separase may not cleave cohesin to promote the metaphase to anaphase transition [30]. However, our results are consistent with the hypothesis that separase is required to cleave whatever remaining cohesin is present on metaphase chromosomes for proper segregation to occur at anaphase onset.

Whether separase has a substrate involved in exocytosis is unknown. However, we find that the protease-dead separase causes cytokinesis failure and inhibits RAB-11 vesicle exocytosis during mitotic cytokinesis. These data are consistent with a model whereby separase cleaves a substrate to promote exocytosis (Fig 9), similar to its function during chromosome segregation and centriole licensing. On its own, SEP-1^PD^::GFP does not cause a severe cytokinesis defect, but synergistically inhibits cytokinesis when *syx-4* is depleted, while chromosome segregation is more obviously defective. It is worth nothing that *C. elegans* centromeres are holocentric [42], meaning that cohesin must be cleaved along the entire chromosome instead of a point centromere as in other organisms and thus chromosome segregation could be more sensitive to delayed cohesin cleavage. Therefore, separase likely cleaves substrates involved in several different process, but the effects imposed by SEP-1^PD^::GFP vary in different events.

**Fig 9.**
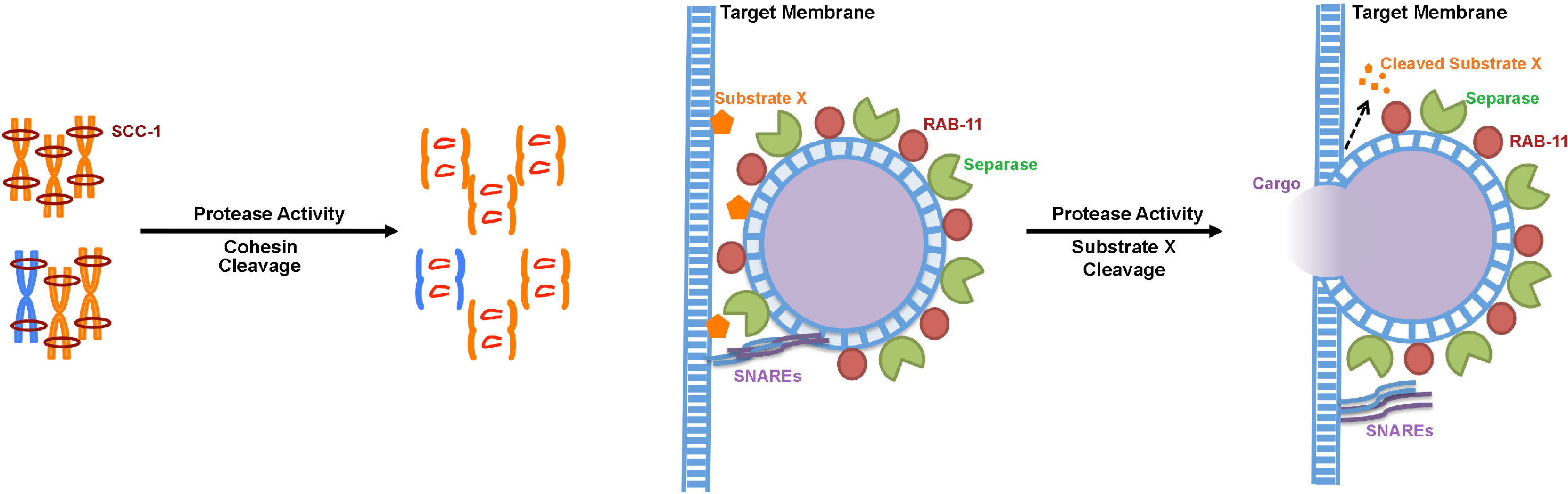
Working model of separase function in exocytosis during cytokinesis. Separase cleaves cohesin kleisin subunit SCC-1 during mitotic anaphase and promotes chromosome segregation. In cytokinesis, separase colocalizes with RAB-11 vesicles. SNAREs including SYX-4 promote vesicle fusion with target membrane. Our results suggest that separase cleaves an unknown substrate to promote exocytosis.

There are several possible explanations for these observations. The first is that our over-expression levels are not high enough to effectively block cleavage of a putative vesicle target, but is sufficient to inhibit chromosome segregation. This could be due to the affinity of separase toward different substrates. While substrate cleavage may be involved in exocytosis, delayed cleavage may not be sufficient on its own to block exocytosis in the presence of all other factors that promote exocytosis. Consistent with this, depletion of separase does not completely block centriole separation and other factors minimize the resulting phenotypes [29]. Certainly the local environment at chromosomes, centrioles and vesicles will be quite different. This could impact how stably separase can interact with substrates and thus how well SEP-1^PD^::GFP can inhibit substrate cleavage. Indeed, separase catalytic activity toward cohesin is much greater in the presence of DNA [43], while the fluid environment of a membrane may not have the same effect. In addition to substrate affinity, SEP-1^PD^::GFP may displace endogenous separase from chromosome more readily than it does in the membrane. Therefore, there may also be differences in the relative amounts of transgenic SEP-1^PD^::GFP to the amount of endogenous separase at different cellular locations. The relative amounts of endogenous vs. transgenic separase protein may also explain why we generally observed less severe meiotic phenotypes vs. mitotic phenotypes. Future studies will be required to resolve these issues.

Our observation show that RAB-11 is recruited to cortical granules much earlier than separase, which suggests an ordered recruitment of regulators to these vesicles prior to their exocytosis in anaphase. Defining the pathway and signals that control the timing sequence of this recruitment process will be important to better understand how the cell cycle and potentially other pathways coordinate vesicle trafficking during cell division. Whether the same process occurs during mitotic cytokinesis will require much better imaging conditions since the individual vesicles are small and dynamic as they move along the spindle. Interestingly, SEP-1^PD^::GFP associates with plasma membrane for an extended period of time after cortical granule exocytosis, however, RAB-11 does not. This indicates that RAB-11 is not required for SEP-1^PD^::GFP to remain associated with the plasma membrane and may not be the substrate of the separase during exocytosis. This result is consistent with previous observations that depletion of RAB-6, but not RAB-11, prevents recruitment of separase to cortical granules [41]. It is still possible that separase may cleave RAB-11 interacting proteins. This could indicate that separase affects a different step in exocytosis than the membrane docking and tethering functions mediated by RAB-11. For example, separase might cleave a substrate that allows vesicles to move forward in the exocytosis pathway, i.e., moving from a docked to a primed state [44]. The timing when cortical granules undergo different steps of exocytosis is unknown, but it is possible that the early steps are completed by the time that separase is completely transferred to vesicles in anaphase. Separase may cleave RAB-11 interacting proteins, such as RAB-11 GEFs, to regulate RAB-11 activity during exocytosis [45]. Another possibility is that separase cleaves an inhibitor of exocytosis, such as the complexin protein that prevents SNAREs from completing vesicle fusion prematurely [46]. Identifying a putative vesicle target that separase cleaves to promote exocytosis is a primary pursuit for future investigation.

While separase is a protease, critical non-proteolytic functions of separase are required for mitotic exit. Previously, three *C. elegans* separase mutant alleles have been identified, all of which map outside of the protease domain [47]. Interestingly, each of these mutants cause defects in cortical granule exocytosis and mitotic cytokinesis failure, but minimal chromosome segregation defects [47]. Furthermore, these mutants are rescued by loss of phosphatase 5 (*pph-5*), which might represent a signaling pathway that controls exocytosis [47]. While our results suggest that separase has a substrate involved in exocytosis, we cannot rule out non-proteolytic functions that may also impact exocytosis. For example, Cdk5 is involved in the regulation of synaptic vesicle exocytosis via phosphorylation of munc18 [48]. Separase may regulate CDK or perhaps another signaling pathway to control exocytosis. Ultimately, separase may have both proteolytic and non-proteolytic functions that collaborate to promote exocytosis during anaphase. This might be required to ensure that separase promotes exocytosis after a significant delay in anaphase, which occurs during both meiosis and mitosis. Elucidating how the precise control of separase function leads to exocytosis during anaphase will be an important goal of future studies.

## Materials and Methods

### *C. elegans* Strains

*C. elegans* strains were maintained with standard protocols. Strains containing the protease-dead SEP-1^PD^::GFP transgene were maintained on *gfp* RNAi feeding plates, then transferred onto regular OP50 bacteria plates according to previous protocols [24]. WH520 hemizygous males were backcrossed to UNC-119 animals to propagate F1 animals for analysis and further propagation. Some strains used in this study were obtained from the Caenorhabditis Genetics Center (CGC). Strain RQ372 was gift from Dr. Risa Kitagawa. JAB18 was created by crossing WH520 males with OD56 hermaphrodites [24]. JAB156 was generated by crossing WH520 males with EKM41 hermaphrodites, and subsequent generations were maintained on *gfp* RNAi. At F2 generation following the cross, L4 stage worms were singled from the original *gfp* RNAi feeding plates. We screened the F3 adults for the presence of PH::mCherry transgenes by microscopy. Then approximately half of the PH::mCherry positive worms at L4 stage were moved to OP50 plates for 3-4 generations, and screened for the presence of both transgenes. The protocol was repeated until double homozygous transgenic lines were obtained, after which the line was maintained on *gfp* RNAi (Table 1).

**Table 1.**
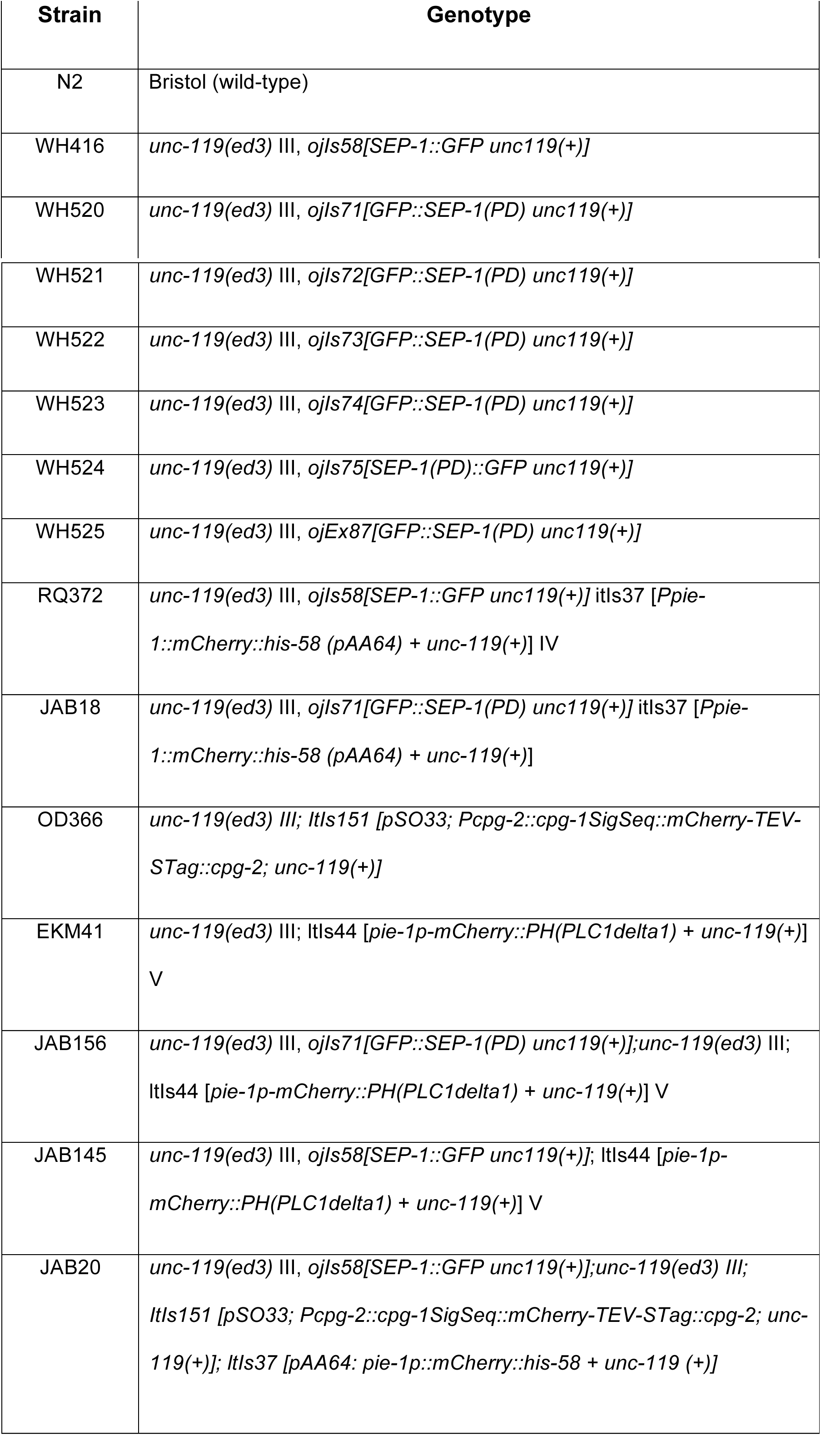
Strains used in this study.

### RNAi Treatment

The *gfp* and *syx-4* RNAi feeding constructs were previously described [24, 33] and *scc-1* RNAi was obtained from the Ahringer library. To silence the target genes, L4 hermaphrodites were picked onto lawns of IPTG-induced RNAi feeding bacteria. In order to provide the optimal RNAi effect for target genes silencing, RNAi cultures were grown till log phase. Then the log phase RNAi bacteria were spread on plates containing NGM agar with 1 mM IPTG and the plates were incubated at 15 °C for 24-48 hours to optimally induce the T7 promoter expression [49]. Worms were grown on RNAi plates at 20°C or 25 °C for the amount of time indicated in the manuscript for different experiments.

### Microscopy

For live imaging, young adult worms were dissected in M9 buffer and embryos were mounted on agar pads as previously described [24]. For imaging of meiotic embryos, or potentially osmotic sensitive embryos, young adults were dissected and mounted in blastomere culture media by hanging drop to relieve mechanical and osmatic pressure [50]. Live cell imaging was performed on a spinning disk confocal system that uses a Nikon Eclipse inverted microscope with a 60 X 1.40NA objective, a CSU-22 spinning disc system and a Photometrics EM-CCD camera from Visitech International. Images were acquired by Metamorph (Molecular Devices) and analyzed by ImageJ/FIJI Bio-Formats plugins (National Institutes of Health) [51, 52].

### Statistics

Quantification of SEP-1::GFP and RAB-11::mCherry at the midbody was performed in Image J by measuring the fluorescence intensity at the midbody in frames with the brightest signal shortly after furrow ingression was completed. Embryos were shifted to 25°C to improve signal, but caused abnormal aggregates of RAB-11::Cherry in some embryos, which were not included in the analysis. To account for variations in imaging and z-depth, we calculated the ratio of the intensity at the midbody relative to cytoplasm. Cytoplasmic signal was determined by averaging the intensities from three separate regions in the same image. Statistical significance was determined by p value from an unpaired two-tailed t-test.

## Acknowledgements

We appreciate the CGC (University of Minnesota) funded by the NIH Office of Research Infrastructure Programs (P40 OD010440) which provided some *C. elegans* strains. We are grateful to members of the Bembenek laboratory, Michael Melesse, Christopher Turpin, Nicolas Mattson, for productive discussion and preparing reagents. We also thank Bruce McKee, Maitreyi Das, Nasser Rusan and Don Fox for critical feedback on the manuscript.

---

S1 Fig. Testing different lines expressing SEP-1^PD^::GFP

(A) Embryonic lethality in F2 broods from hemizygous SEP-1^PD^::GFP/- transgenic hermaphrodites from backcrossing propagation strategy. The number of generations the strain was backcrossed is also indicated. (B) Embryonic lethality of homozygous SEP-1^PD^::GFP lines propagated off *gfp(RNAi)* for 5-6 generations. Each data with error bars represents the average of embryonic lethality from 6-10 singled worms (n=singled worm number: total embryo count).

S2 Fig. SEP-1^PD^::GFP does not affect global cell cycle timing during mitosis

Different stages of mitosis: (A) Nuclear envelope intact in prophase, (B) Nuclear envelope breakdown (NEBD), indicated by separase in the nucleus and H2B::Cherry nucleoplasmic signal dispersing in prometaphase, (C) Chromosome alignment in metaphase, and (D) Initiation of furrow ingression as observed by DIC acquired simultaneously with the fluorescent images. (E) Timing from NEBD to metaphase in SEP-1^WT^::GFP and SEP-1^PD^::GFP embryos with and without *scc-1(RNAi)*. F. Timing from NEBD to furrow ingression in SEP-1^WT^::GFP and homozygous SEP-1^PD^::GFP embryos with and without *scc-1(RNAi)*.

S3 Fig. SEP-1^PD^::GFP affects furrow ingression (SEP-1^PD^::GFP reduces the rate of furrow ingression)

(A) Kymograph depicting the timing of furrow ingression in AIR-2::GFP (green) expressing PH::mCherry and H2B::mCherry (red) after furrow initiation (upper kymograph). Middle kymograph shows only the red channel (PH::mCherry and H2B::mCherry) in the AIR2::GFP embryo. The lower kymograph shows the delayed cleavage furrow (PH::mCherry channel) in a homozygous SEP-1^PD^::GFP embryo. (B) Quantification of the furrow ingression rate in different genotypes, depletion of *scc-1* does not rescue the slower furrow ingression in SEP-1^PD^::GFP (p=0.29).

S4 Fig. Depletion of t-SNARE *syx-4* enhances cytokinesis failure in hemizygous SEP-1^PD^::GFP/-; RAB-11::mCherry embryos.

(A-C) Representative images of mitotic cytokinesis failure in hemizygous SEP-1^PD^::GFP/- (green) embryos expressing RAB-11::mCherry (red) and treated with *syx-4(RNAi)*. (D) The percentage of embryos displaying cytokinesis failure in indicated conditions.

S5 Fig. SEP-1^PD^::GFP does not affect RAB-6 trafficking during cytokinesis

(A) Kymograph of the furrow region showing that RAB-6::mCherry (red) and SEP-1^WT^::GFP (green) do not accumulate in the furrow. (B) Accumulation of hemizygous SEP-1^PD^::GFP/- (green) is observed at the furrow and midbody, but not RAB-6::mCherry (red).

S6 Fig. Both SEP-1^WT^::GFP and SEP-1^PD^::GFP/-Co-localized with cortical granule marker CPG-2::mCherry during cortical granule exocytosis.

Both SEP-1^WT^::GFP (green) and SEP-1^PD^::GFP/- (green) colocalize with CPG-2::mCherry (red) at cortical granules. Insets are magnified images of CGs showing that CPG-2, a cargo protein that should be in the lumen of the vesicle, appears to be surrounded by separase signal, which is likely associated with the vesicle membrane. Scale Bars, 10 μm.

Supplemental Movie 1. Chromosome segregation during mitosis

Examples of embryos expressing H2B::mCherry (red) with SEP-1^WT^::GFP (green, left embryo) or homozygous SEP-1^PD^::GFP (green) with slightly lagging (middle) or bridging (right embryo) chromosome segregation defects during the first mitotic division. Bottom movies show H2B::mCherry only to display bridges. Images were acquired in a single z plane every 6 seconds. Playback rate is 10 frames/ second.

Supplemental Movie 2. Chromosome segregation during meiosis I

Embryos expressing SEP-1^WT^::GFP (green, top left) or homozygous SEP-1^PD^::GFP (green, top right) and H2B::mCherry (red) during meiosis I. Bottom movies show H2B::mCherry only to display bridges. Images were acquired in a single z plane every second. Playback rate is 10 frames/ second.

Supplemental Movie 3. Cytokinesis failure of SEP-1^PD^::GFP is synergistically enhanced by *syx-4(RNAi)*

Cytokinesis failures in *syx-4 (RNAi)* embryos expressing PH::mCherry (red) and SEP-1^WT^::GFP (green, left) is rare, but homozygous SEP-1^PD^::GFP (green, right) shows a high incidence of cytokinesis failure. Images were acquired in a single z plane every 6 seconds. Playback rate is 10 frames/ second.

Supplemental Movie 4. Depletion of t-SNARE *syx-4* caused cytokinesis failure in hemizygous SEP-1^PD^::GFP/-; RAB11::mCherry expression embryos.

Embryo expressing hemizygous SEP-1^PD^::GFP/- (green) and RAB-11::mCherry (red) with *syx-4 (RNAi)* treatment fails cytokinesis during first mitotic cell division. Images were acquired in a single z plane every 6 seconds. Playback rate is 10 frames/ second.

Supplemental Movie 5. SEP-1^PD^::GFP/- affects RAB-11 trafficking during cytokinesis.

Embryos expressing RAB-11::mCherry (red) and SEP-1^WT^::GFP (green, upper movies) or hemizygous SEP-1^PD^::GFP/- (green, lower movies) with control (left) or *syx-4 (RNAi)* (right) treatment. Images were acquired in a single z plane every 6 seconds. Playback rate is 10 frames/ second.

Supplemental Movie 6. Separase localization during cortical granule exocytosis

Embryos expressing both SEP-1^WT^::GFP (green, left) or hemizygous SEP-1^PD^::GFP/- (green, right) co-localized with CPG-2::mCherry (red) on the cortical granules. Note that CPG-2::mCherry only localizes to a subset of separase labeled vesicles. Images were acquired in a single z plane every 6 seconds. Playback rate is 10 frames/ second.

Supplemental Movie 7. SEP-1^PD^::GFP delays completion of cortical granule exocytosis

Embryos expressing both SEP-1^WT^::GFP (green, left) or hemizygous SEP-1^PD^::GFP/- (green, right) with H2B::mCherry (red). In homozygous SEP-1^PD^::GFP embryos, cortical granule exocytosis is delayed compared with SEP-1^WT^::GFP. SEP-1^PD^::GFP associates with plasma membrane much longer than WT after CGE. Images were acquired in a single z plane every 3 seconds. Playback rate is 10 frames/ second.

Supplemental Movie 8. Separase associates with plasma membrane after CGE.

SEP-1^WT^::GFP (green, left) and homozygous SEP-1^PD^::GFP (green, right) co-localized with plasma membrane marker PH::mCherry (red) after CGE. SEP-1^PD^::GFP associated with plasma membrane longer compared with SEP-1^WT^::GFP. Images were acquired in a single z plane every 6 seconds. Playback rate is 10 frames/ second.

Supplemental Movie 9. Separase regulates cortical granules exocytosis as well as RAB-11.

Embryos expressing RAB-11::mCherry (red) with SEP-1^WT^::GFP (green, left) and hemizygous SEP-1^PD^::GFP/- (green, right) during anaphase I. RAB-11 co-localizes on the same population of vesicles with both forms of separase. After exocytosis, SEP-1^PD^::GFP/- associates with the plasma membrane, while RAB-11::mCherry rapidly disappears. Images were acquired in a single z plane every 6 seconds. Playback rate is 10 frames/ second.

